# *Thm2* interacts with paralog, *Thm1*, and sensitizes to Hedgehog signaling in postnatal skeletogenesis

**DOI:** 10.1101/2020.01.26.920165

**Authors:** Bailey A Allard, Wei Wang, Tana S Pottorf, Hammad Mumtaz, Luciane M Silva, Damon T Jacobs, Jinxi Wang, Erin E Bumann, Pamela V Tran

## Abstract

Ciliopathies are genetic syndromes that link osteochondrodysplasias to dysfunction of primary cilia. Primary cilia extend from the surface of bone and cartilage cells, to receive extracellular cues and mediate signaling pathways. Mutations in several genes that encode components of the intraflagellar transport-A ciliary protein complex have been identified in skeletal ciliopathies, including *THM1*. Here, we report a role for genetic interaction between *Thm1* and its paralog, *Thm2,* in skeletogenesis. THM2 localizes to the ciliary axoneme, but unlike its paralog, *Thm2* deficiency does not affect ciliogenesis and *Thm2-*null mice survive into adulthood. Since paralogs often have redundant functions, we crossed a *Thm1* null (*aln*) allele into the *Thm2* colony. After 5 generations of backcrossing the colony onto a C57BL6/J background, we observed that by postnatal day 14, *Thm2^-/-^; Thm1^aln/+^* mice are smaller than control littermates. *Thm2^-/-^; Thm1^aln/+^* mice exhibit shortened long bones, narrow ribcage, shortened cranium and mandibular defects. Mutant mice also show aberrant architecture of the tibial growth plate, with an expanded proliferation zone and diminished hypertrophic zone, indicating impaired chondrocyte differentiation. Using microcomputed tomography, *Thm2^-/-^; Thm1^aln/+^* tibia were revealed to have reduced cortical and trabecular bone mineral density. Deletion of one allele of *Gli2*, a major transcriptional activator of the Hedgehog (Hh) pathway, exacerbated the small phenotype of *Thm2^-/-^; Thm1^aln/+^* mice and caused small stature in *Thm2*-null mice. Together, these data reveal *Thm2* as a novel locus that sensitizes to Hh signaling in skeletal development. Further, *Thm2^-/-^; Thm1^aln/+^* mice present a new postnatal ciliopathy model of osteochondrodysplasia.

## Introduction

Skeletal dysplasias affect 1:5,000 births and range in severity from perinatal lethality to craniofacial dysmorphogenesis and short stature. These disorders are often heritable, and identifying the causative mutations has been instrumental in improving diagnosis, understanding mode of inheritance, and identifying therapeutic targets.

Conversely, genetic disorders that cause skeletal dysplasias provide valuable insight into underlying molecular and cellular mechanisms. One such class of genetic disorders are ciliopathies, which result from mutation and dysfunction of primary cilia. The primary cilium is a solitary, sensory organelle that protrudes from the surface of most mammalian cells, including bone and cartilage cells^1, 2^. As such, ciliopathies affect multiple organ systems, and a proportion of ciliopathies manifest osteochondrodysplasias^3, 4^. These skeletal ciliopathies include Short-Rib Polydactyly Syndromes (I-IV), Jeune Syndrome, Oro-facial-digital syndrome type 1, Ellis van Creveld syndrome, and Sensenbrenner Syndrome, which manifest to varying degrees, shortened long bones, narrow rib cage, polydactyly and craniofacial defects. Fibrocystic diseases of the kidney, liver and pancreas can also arise. These ciliopathies reveal that primary cilia are potent modifiers of skeletal growth.

Primary cilia detect and transduce extracellular mechanical and chemical cues into signaling cascades to ultimately affect cellular behavior. These antenna-like organelles are dynamic structures, formed and maintained by intraflagellar transport (IFT), which is the bi-directional transport of cargo (structural and signaling proteins) by multiprotein complexes along a microtubular axoneme^5, 6^. IFT-B proteins, together with the kinesin motor, transport cargo from the base to the tip of the cilium in anterograde IFT, while IFT-A proteins and the dynein motor return proteins from the ciliary tip to the base in retrograde IFT. IFT-A proteins also mediate ciliary entry of membrane-associated and signaling proteins^7–10^. Primary cilia are required to transduce the Hedgehog (Hh) signal^11^. The presence of Hh ligand induces ciliary enrichment of the Smoothened signal transducer and culminates in activation of the *Gli* transcription factors, of which GLI2 is the primary transcriptional activator^12, 13^. The differing roles of IFT-B versus -A can result in contrasting ciliary and Hh signaling phenotypes in *Ift* mutant mouse embryos. Loss of Ift-B or kinesin, generally results in loss of cilia and Hh activity, and mid-gestational lethality^14–16^, while deletion of Ift-A causes shortened primary cilia with protein accumulation in a bulbous distal tip. Further, two Ift-A mouse embryonic mutants show overactivation of the Hh pathway, and perinatal lethality^17, 18^. Yet in contrast to these earlier reports, deletion of kinesin subunit, *Kif3a*, in neural crest cells increased GLI activity in the frontonasal prominence of the developing face^19^, while deletion of Ift-A gene, *Thm1*, in glial cells decreased Hh signaling in the developing cerebellum^20^. Thus, whether an IFT protein positively or negatively regulates Hh or GLI activity is cell-specific and/or context-dependent.

In patients with skeletal ciliopathies, mutations have been identified in all of the 6 characterized IFT-A components (*IFT43*^21, 22^*, IFT121*^22^*, IFT122*^23, 24^*, WDR19/IFT144*^25^*, IFT140*^26^, and *THM1*^27^) and in only 3 of the 14 IFT-B components (*IFT80*^28^, *IFT52*^29, 30^ and *IFT172*^31^). In accordance with the human genetics studies, all reported IFT-A mouse mutants have exhibited polydactyly, and abnormal development of the thoracic ribs during embryogenesis^32–34^. Additionally, *Ift80*-hypomorphic mice that survive past birth, and mice with conditional deletion of IFT-B gene, *Ift88,* in the limb bud and craniofacial mesenchyme, exhibit polydactyly, shortened long bones, and disorganization of the growth plate^35, 36^. Thus, both IFT-B and -A are essential for mammalian skeletal growth, although the human genetics studies may suggest that IFT-A mutations are more compatible with life.

Previously, we identified *THM1* (TPR-containing Hh modulator 1; also known as *TTC21B*) as an IFT-A component and regulator of Hh signaling^17^. A germ-line mutation resulting in THM1 loss in mouse causes shortened long bones, split and fused ribs, and polydacytyly. *THM1* mutations have also been identified in patients with Jeune Syndrome (JATD)^37^, which is characterized by shortened long bones, craniofacial defects, and a narrow rib cage that often inhibits lung development and causes 60-80% mortality among infants and children^38^. We also identified a paralog of *THM1*, which we called *THM2* (also known as *TTC21A*)^17^. THM1 and THM2 are orthologues of the IFT-A protein, IFT139, in *Chlamydomonas reinhardtii*, and have similar levels of homology to *IFT139/FLA16. Thm1* and *Thm2* have similar predicted protein structures with multiple TPR domains and similar RNA expression patterns during mouse development. A recent report revealed that *THM2* is required for male fertility in humans and mice^39^. However, a role for *Thm2* in early postnatal development has not been elucidated. Here by generating *in vitro* and *in vivo* models of *Thm2* deficiency and implementing genetic crosses, we aimed to establish the subcellular localization of THM2 and its role in postnatal skeletogenesis.

## Results

### THM2 localizes to primary cilia

To characterize the subcellular localization of THM2, we had a custom-made antibody that recognizes a unique amino acid sequence in exon 22 of THM2 generated (Proteintech). To determine the specificity of the antibody, we generated human embryonic kidney (293T) *THM2* knock-down (kd) clonal cell lines using lentiviruses expressing *THM2* shRNA. Western blot analysis revealed *THM2* kd, clone 2-1, showed the most effective knockdown (Figure 1B). We analyzed cilia length in EV 2-5 (empty vector control) and *Thm2* kd 2-1 clonal cell lines by immunostaining for ARL13B, a ciliary membrane marker (Figure 1C). Quantification of ciliary lengths demonstrated similar lengths between EV and *THM2* kd cell lines (Figure 1D), revealing THM2 deficiency does not affect ciliogenesis. However, immunostaining of retinal pigment epithelial (RPE) cells for THM2 showed co-localization with acetylated *α*-tubulin, a marker of the ciliary axoneme (Figure 1A), indicating that THM2 localizes to primary cilia.

**Figure 1.**
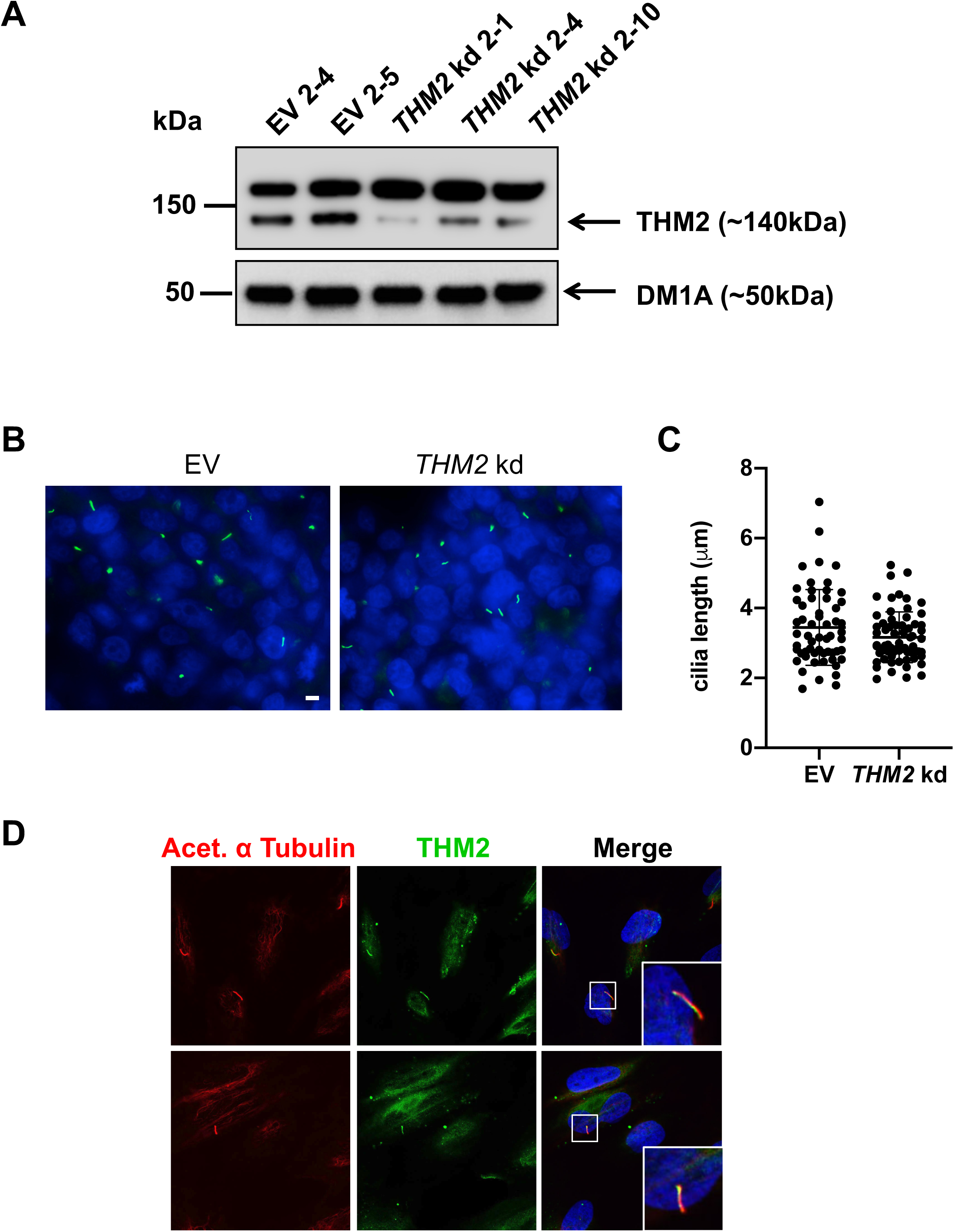
THM2 localizes to the primary cilium, but is dispensable for ciliogenesis. A) Western blot for THM2 on cellular extracts of 293T EV (control) and *THM2* kd clonal lines. B) Immunostaining for ARL13B (green) in 293T EV and *THM2* kd (clone 2-1) cells. C) Quantification of cilia length. Scale bar - 10µm. Each data point represents an individual cilium. Graphs represent mean ± SD. D) Immunostaining for THM2 (green) and acetylated α-tubulin (red) in RPE cells. Scale bar equals 5µm.

### Generation of Thm2 knockout mouse model

Using a *Thm2* knock-out construct from the Knockout Mouse Project (KOMP) repository, we generated a *Thm2* null allele, in which exon 6 was excised, creating a premature stop codon (Figure 2A). Primers flanking exon 6 were designed to amplify a WT band of 1,086bp and a recombined band at approximately 400bp (Figure 2B). Additionally, since this PCR reaction can favor the amplification of the smaller recombined (*Thm2-null)* allele, we designed additional primers to amplify a 461bp product of the WT allele only (Figure 2B). In accordance with amplification of the mutant band, Western blot analysis confirmed the loss of THM2 protein in testis protein extracts of a *Thm2-null* mouse (Figure 2C).

**Figure 2.**
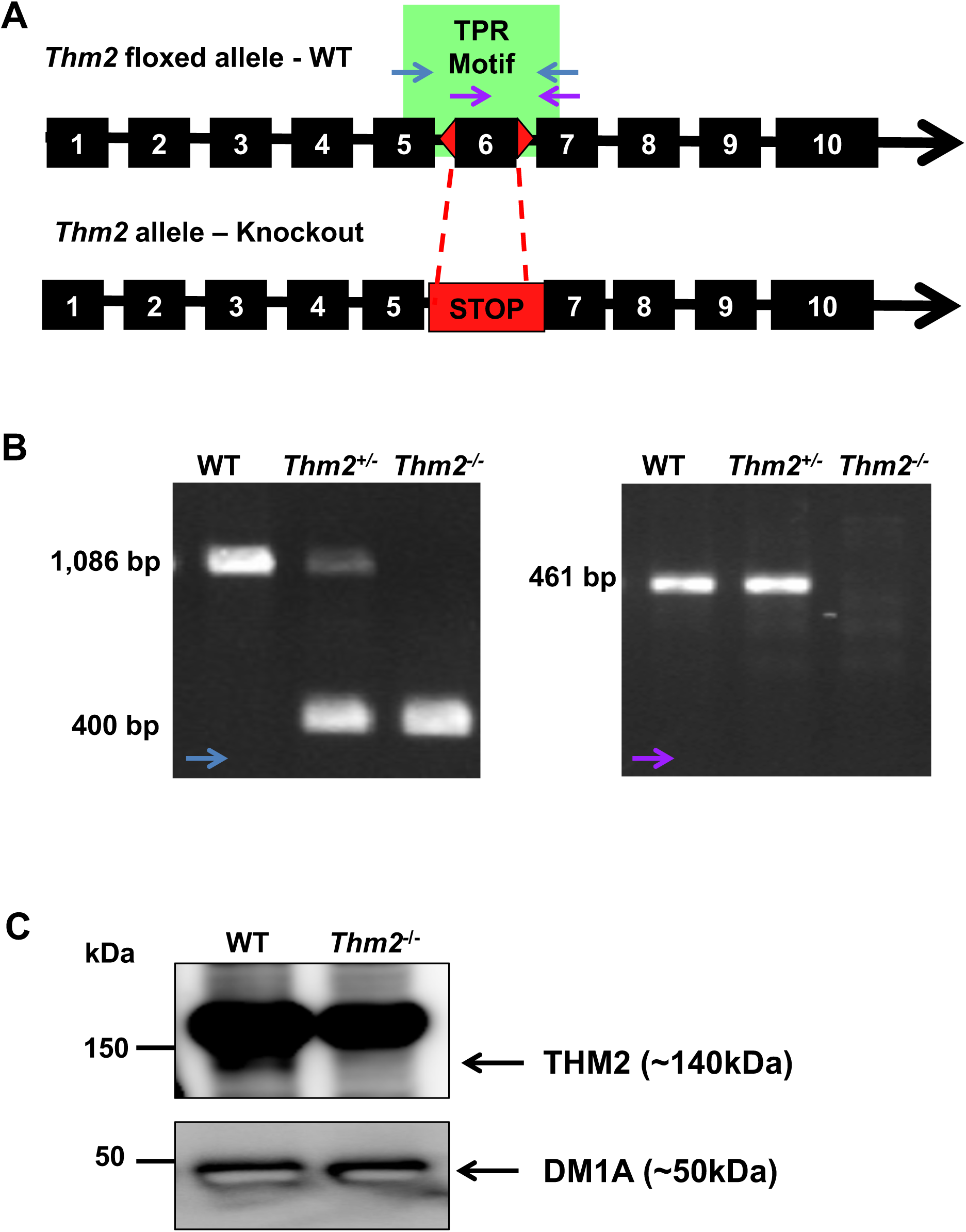
*Thm2* knock-out mouse was generated using KOMP construct. A) Schematic diagram of the KOMP construct in genomic DNA of *Thm2*. Exons are represented by black boxes and introns by the black line. Lox P sites surrounding exon 6 are depicted as red triangles. Genomic primers are indicated in blue (WT and recombination band) and purple (WT band). Splicing of exon 6 creates a premature stop codon. B) PCR analysis of WT, *Thm2^+/-^*, and *Thm2^-/-^* genotypes using primers indicated in blue and additional confirmation of the WT allele in WT and *Thm2^+/-^* mice using primers indicated in purple. C) Western blot analysis of P14 testis extracts confirms loss of THM2 in *Thm2* ko mice.

### Thm2^-/-^; Thm1^aln/+^ mice are smaller than control littermates

Unlike most ciliary gene mouse knock-outs, including *Thm1*, which are embryonic lethal^17, 18, 33, 34^, *Thm2*-null mice are born, phenotypically indistinguishable from their littermates, and reach adulthood with seemingly normal health. Since paralogs can have redundant functions^40, 41^, we introduced a mutant allele of *Thm1 (aln)* onto the *Thm2*-null background. The *aln* allele harbors a missense mutation that results in absence of protein^17^. We also backcrossed the colony five generations onto a C57BL6/J background. At P14, *Thm2^-/-^; Thm1^aln/+^* mice were noticeably smaller than their control littermates with reduced body weight and shorter crown-to-rump length (Figures 3A-B). This phenotype was also observed at P21 (Figure 3C), and we noted that a proportion of triple allele mutant mice (7/25 = 28%) do not survive to weaning age.

**Figure 3.**
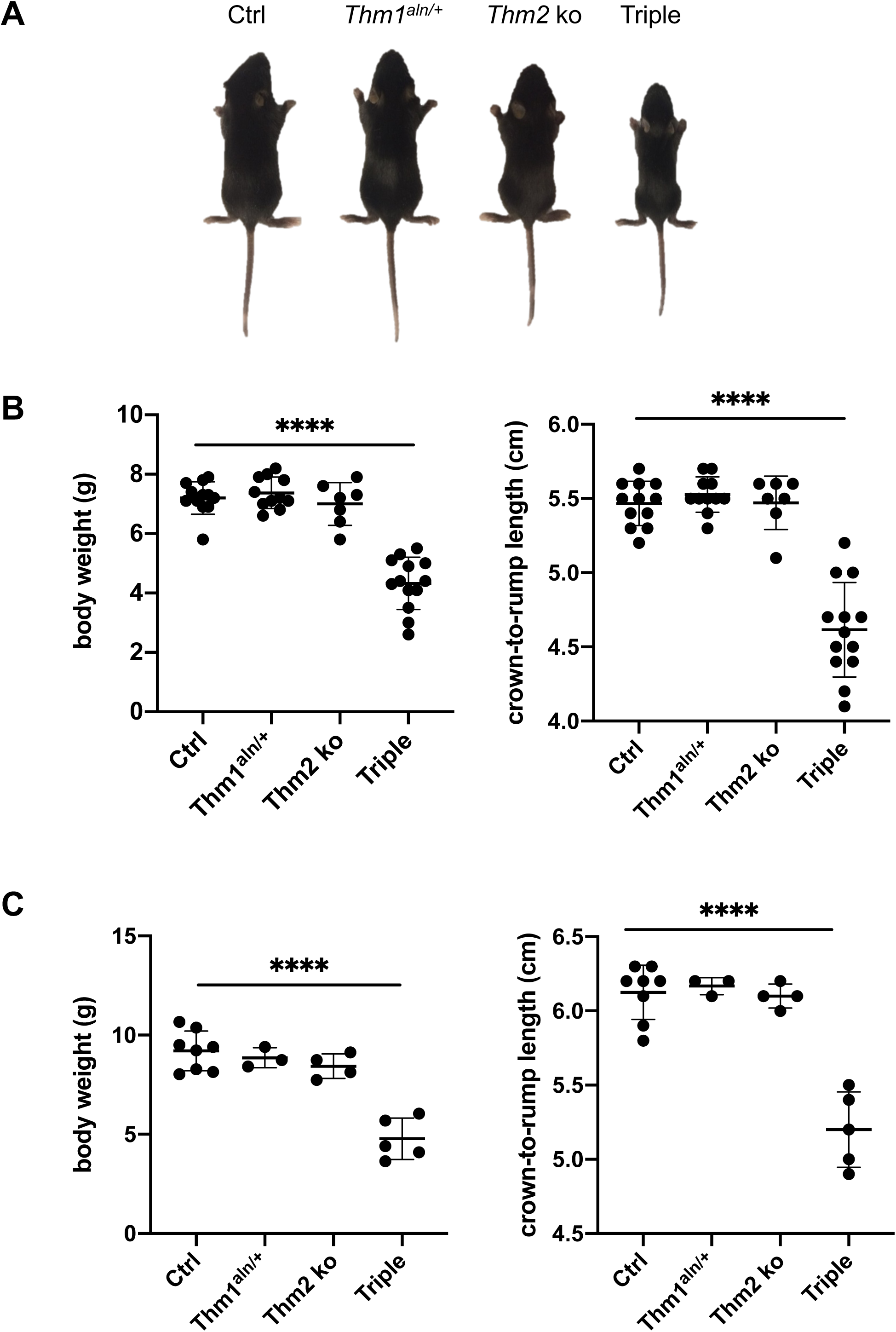
*Thm2^-/-^; Thm1^aln/+^* mice exhibit a runted phenotype. Following 5 generations of backcrossing onto a C57BL6/J background, triple allele mutants showed a runted phenotype. A) Images of mice at P14. Body weights and crown-to-rump lengths (B and C) at P14 and (D and E) P21. Error bars represent mean ± standard deviation. Statistical significance was determined using one-way ANOVA and Tukey’s test. ****p≤0.0001

### Thm2^-/-^; Thm1^aln/+^ lung weight/body weight ratio is increased

Since renal cysts are among the most common clinical manifestations of ciliopathies^42^, we analyzed the kidneys of P14 *Thm2^-/-^; Thm1^aln/+^* mice. Histological analysis revealed normal kidney morphology of *Thm2^-/-^; Thm1^aln/+^* mice (Figure 4A). Total kidney weights were smaller in *Thm2^-/-^; Thm1^aln/+^* mice, but this was proportional to body size since kidney weight/body weight ratios of *Thm2^-/-^; Thm1^aln/+^* mice were similar to those of control littermates (Figure 4A). Similarly, heart weight was lower in triple allele mutant mice, but heart weight/body weight ratios were comparable between control and *Thm2^-/-^; Thm1^aln/+^* mice (Figure 4B). In contrast, although lung weight was lower in triple allele mutant mice, lung weight/body weight ratio was elevated in triple allele mutant mice relative to control mice (Figure 4C).

**Figure 4.**
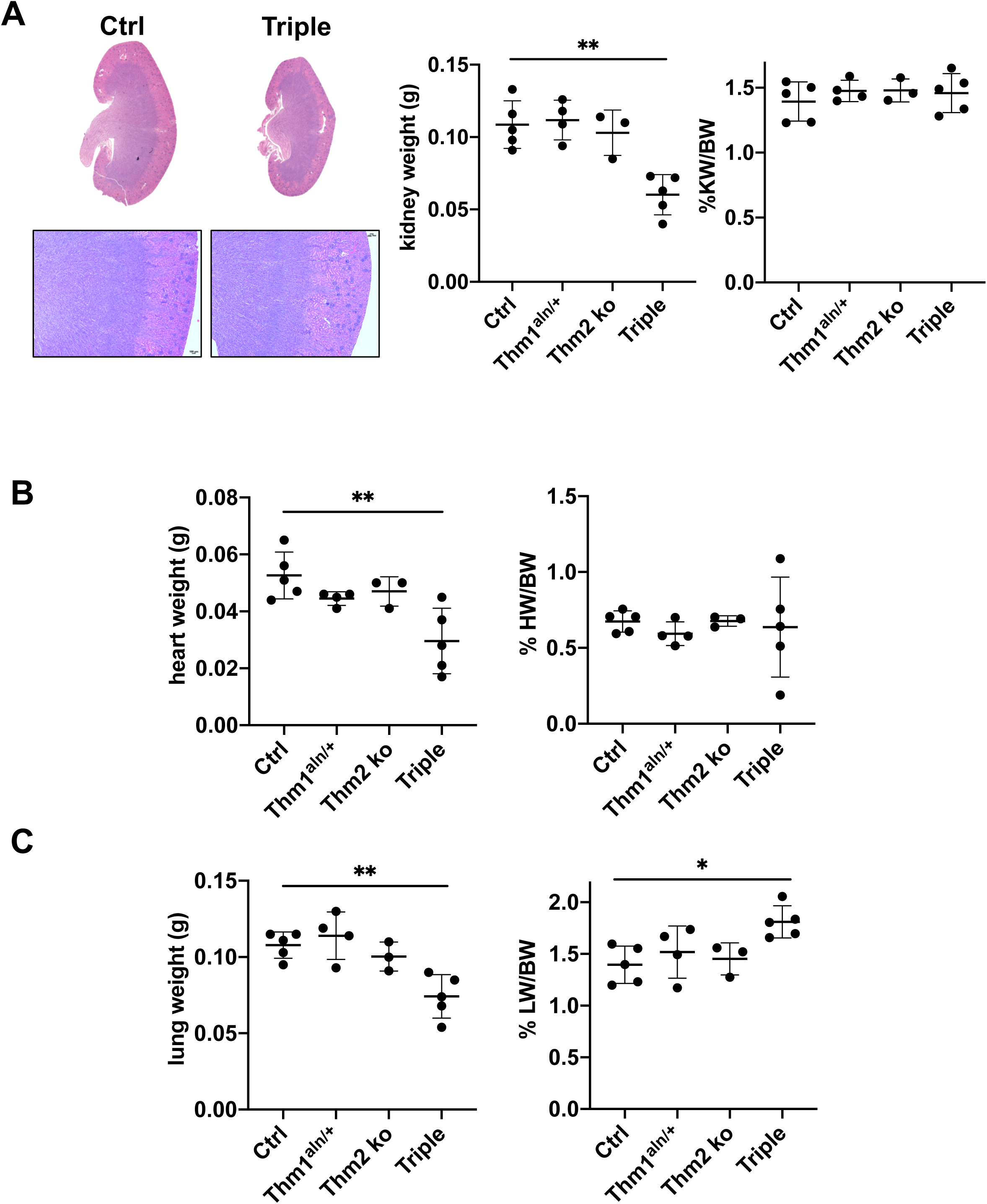
*Thm2^-/-^; Thm1^aln/+^* mice have increased lung/body weight ratios. A) Haematoxylin and eosin staining of kidney sections from P14 mice. Two kidney weights and % kidney weight/body weight ratios. B) Lung weights and % lung weight/body weight ratios. C) Heart weights and % heart weight/body weight ratios. Graphs represent mean ± standard deviation. Statistical significance was determined using one-way ANOVA and Tukey’s test. **p≤0.01, ***p≤0.001

### Thm2^-/-^;Thm1^aln/+^ mice exhibit shortened and less dense bone

To characterize the skeletal phenotype of *Thm2^-/-^; Thm1^aln/+^* mice, we performed skeletal preparations at P14 (Figure 5A), and measured bone lengths. At P14, *Thm2^-/-^; Thm1^aln/+^* mice exhibited shorter long bones, including tibia, femur, humerus, radius and ulna (Figure 5B). Further, the sternum and ribcage diameter of *Thm2^-/-^; Thm1^aln/+^* mice were smaller than those of control littermates. Cranial length, as measured from tip of nose to back of head, was also shorter in *Thm2^-/-^; Thm1^aln/+^* mice. To determine if these reduced bone lengths reflect overall dwarfing of the tripe allele mutants, we examined bone length/crown-to-rump length ratios. Long bone, sternum and ribcage/crown-to-rump ratios were similar between control and triple allele mutants, but the skull/crown-to-rump ratio was slightly increased in triple allele mutants (Figure 5C). Moreover, triple allele mutants showed small mandible/skull ratios, indicative of craniofacial defects (Figure 5D).

**Figure 5.**
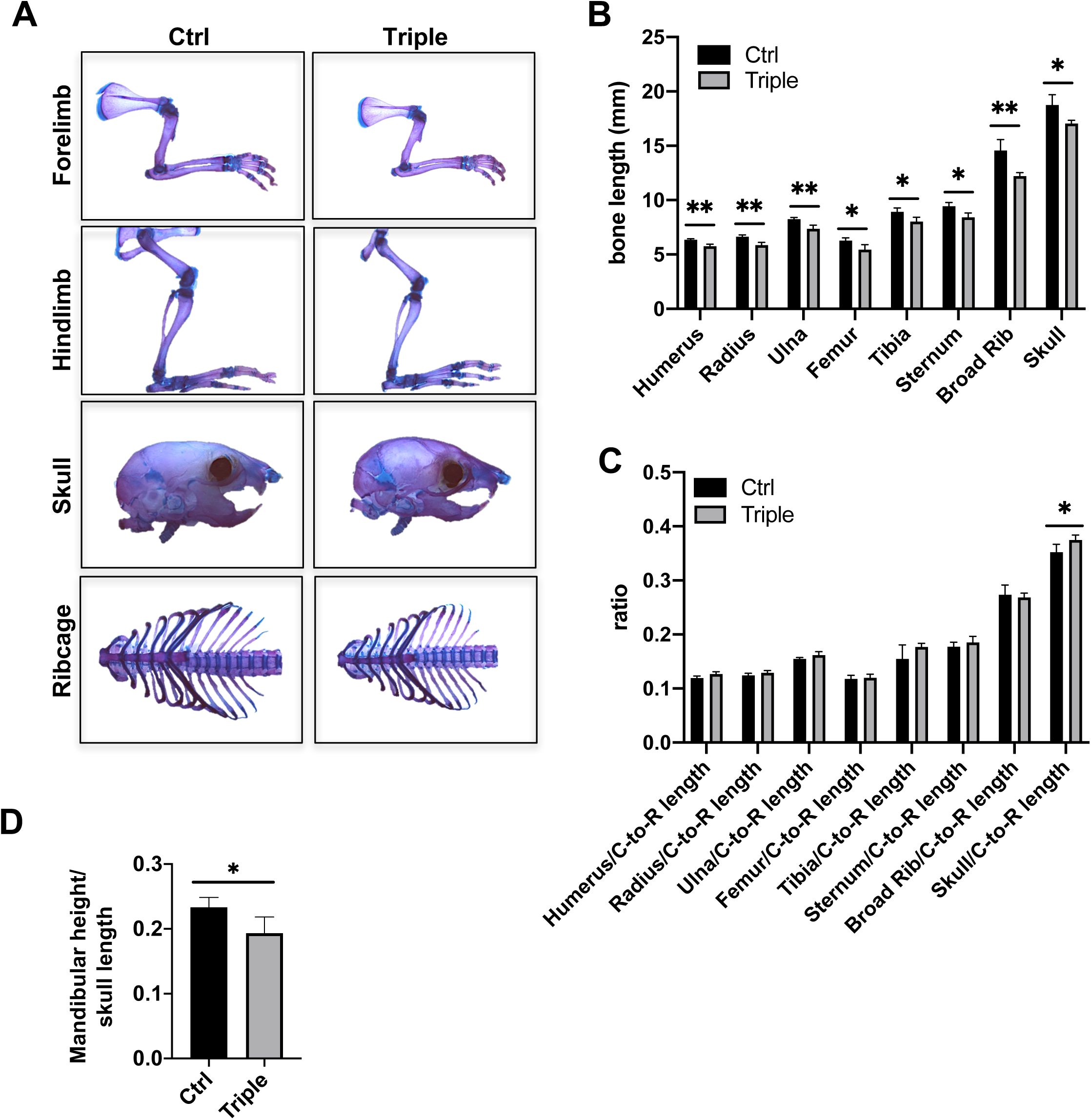
*Thm2^-/-^; Thm1^aln/+^* mice have shorter bones. A) Skeletal preparations at P14 using alizarin red (bone) and alcian blue staining (cartilage). B) Bone lengths. C) Bone/Crown-to-Rump (C-to-R) length ratios. D) Mandibular height/cranial length. Graphs represent mean ± standard deviation of n=4 ctrl and n=4 triple allele mutant mice. Statistical significance was determined using two-tailed, unpaired t-test. *p≤0.05; **p≤0.005

**Figure 6.**
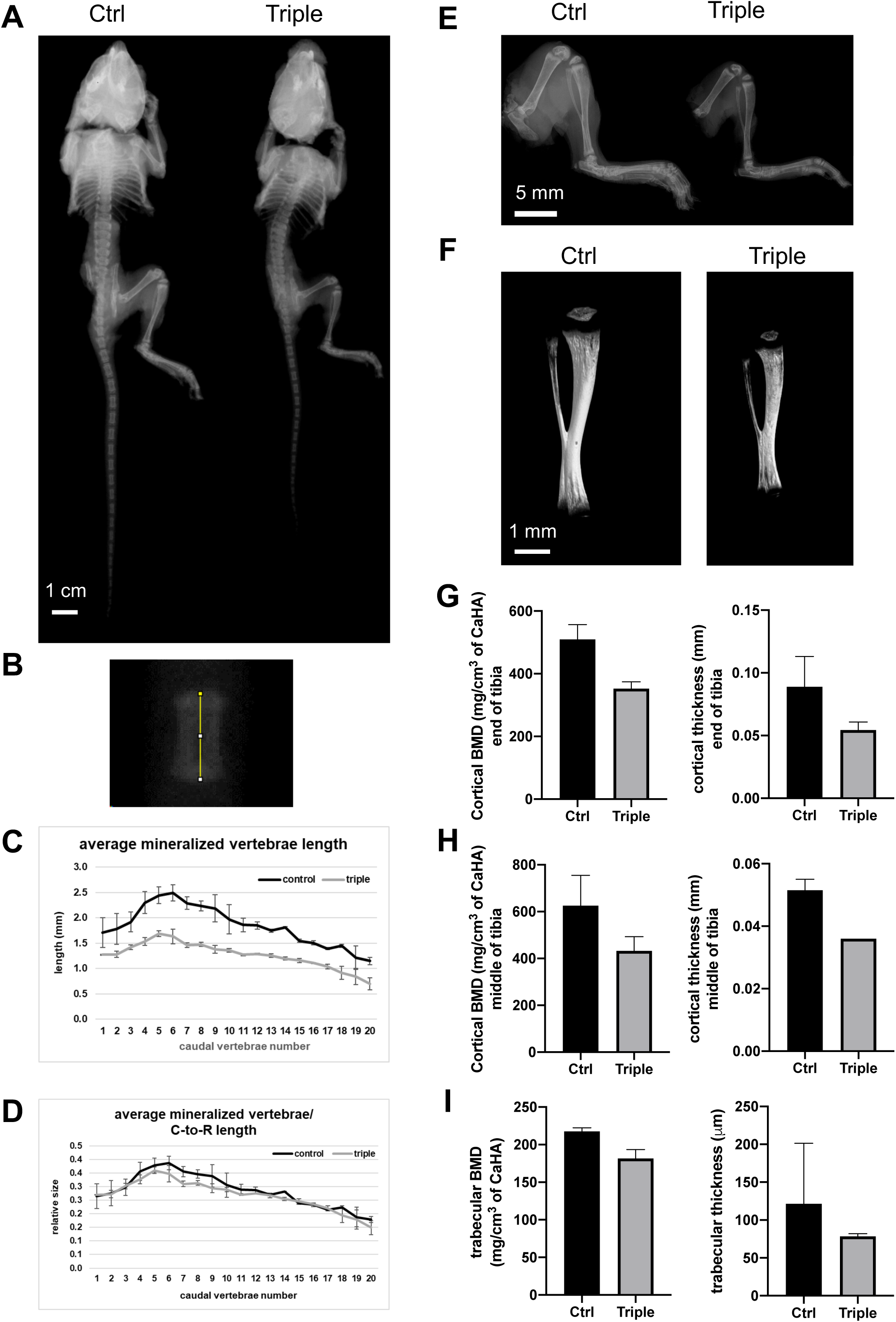
*Thm2^-/-^; Thm1^aln/+^* tibia have less dense bone. Radiographs of A) entire skeleton and of B) vertebra at P14. C) Caudal vertebrae lengths by number. D) Caudal vertebrae length/Crown-to-Rump (C-to-R) length ratios. Graphs represent mean ± standard deviation of n=2 ctrl and n=2 triple allele mutant mice. E) Radiographs of hindlimbs and F) microCT imaging of tibias. G) Cortical bone mineral density (BMD) and thickness at proximal end of tibia. H) Cortical BMD and thickness at mid-diaphysis. I) Trabecular BMD and thickness at proximal end of tibia.

Using radiographs, we further analyzed the *Thm2^-/-^; Thm1^aln/+^* skeleton (Figure 6A and 6B). Individual caudal vertebra length was decreased in *Thm2^-/-^; Thm1^aln/+^* mice (Figures 6C), although proportional to crown-to-rump length (Figure 6D). We also noted that the hindlimb and tibia of *Thm2^-/-^; Thm1^aln/+^* mice were less opaque than those of control mice by radiograph and micro-computed topography (Figures 6E and 6F). This was particularly noticeable when tibias were rotated (Movies S1 and S2). Consistent with this, *Thm2^-/-^; Thm1^aln/+^* tibia showed reduced cortical and trabecular bone mineral density (BMD) and thickness (Figures 6G-6I). Other cortical and trabecular bone properties of control and *Thm2^-/-^; Thm1^aln/+^* tibia were similar (Figure S1). These data demonstrate that *Thm2*, together with *Thm1,* is required for proper postnatal skeletal development.

### Growth plates of Thm2^-/-^;Thm1^aln/+^ show aberrant architecture

We next examined chondrocyte organization in the tibial growth plate at P14. Broadly, the growth plate can be divided into two major regions: the proliferative zone and the hypertrophic zone^43^. As part of endochondral ossification, chondrocyte differentiation occurs. Early chondroblasts are stimulated to proliferate, then become pre-hypertrophic, hypertrophic, and finally undergo cell death and calcification of cartilage. Capillaries and osteoblasts then invade the cartilage matrix and form new bone which replaces cartilage. H&E staining, as well as Safranin O and fast green staining, to mark the cartilage and bone, respectively, of *Thm2^-/-^; Thm1^aln/+^* tibial growth plates revealed an expanded proliferation zone and a reduced hypertrophic zone (Figure 7). We also observed a more basophilic H&E staining and more intense safranin-O staining in the *Thm2^-/-^; Thm1^aln/+^*growth plate cartilage proliferation zone. This may indicate a change in proteoglycan content of the extracellular matrix. Altogether, these defects suggest impaired chondrocyte hypertrophy and endochondral ossification, resulting in shortened long bones seen in *Thm2^-/-^;Thm1^aln/+^ mice*.

**Figure 7.**
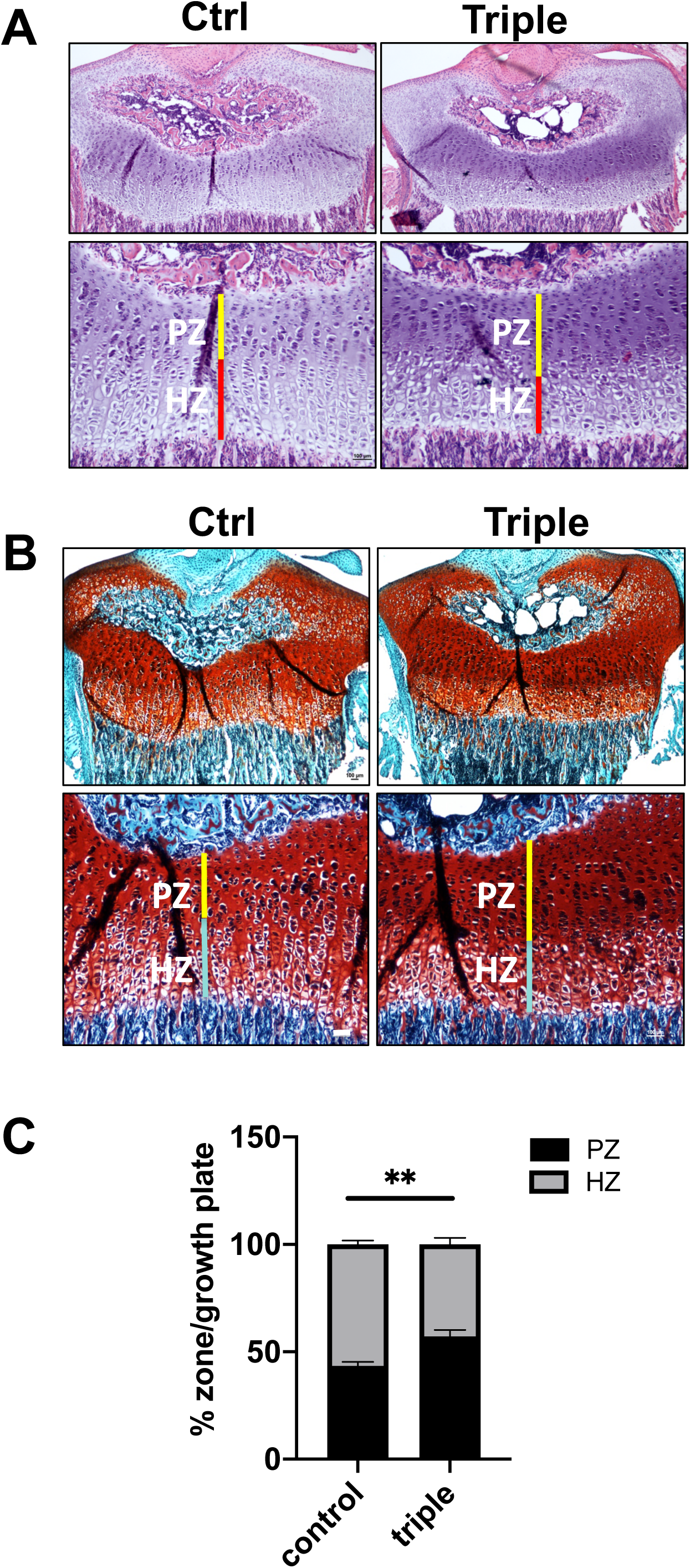
*Thm2^-/-^; Thm1^aln/+^* mice have altered tibial growth plates. A) Haematoxylin and eosin (H&E) staining, and B) Safranin O and Fast Green staining of growth plates at P14. C) Quantification of % proliferation zone (PZ)/growth plate and % hypertrophic zone (HZ)/growth plate from H&E images. Error bars represent SD of n=3 ctrl and n=3 triple allele mutant growth plates. Statistical significance was determined using two-tailed, unpaired t-test. **p≤0.005

### Thm2^-/-^; Thm1^aln/+^cilia length is not affected

While *Thm2* deficiency does not alter cilia length (Figure 1B), we examined whether the additional loss of one allele of *Thm1* affects cilia length. Immunostaining for the ciliary axoneme (acetylated α-tubulin) on chondrocytes of the growth plate showed similar lengths between control and *Thm2^-/-^; Thm1^aln/+^* cilia (Figure 8). Similarly, immunostaining of mouse embryonic fibroblasts (MEF) for ARL13B and γ-tubulin also revealed similar cilia lengths for control, *Thm2-/-,* and *Thm2^-/-^; Thm1^aln/+^* cells, in contrast to *Thm1^aln/aln^* cells, which have shorter cilia length (Figure S1)^44^. While *Thm2^-/-^; Thm1^aln/+^* cilia length is not affected, the altered growth plate architecture and the skeletal phenotype suggest cilia function is disrupted. Similarly, *Ift80*-hypomorphic mutant mice show normal cilia structure, but impaired Hh signaling and a postnatal skeletal phenotype^35^.

**Figure 8.**
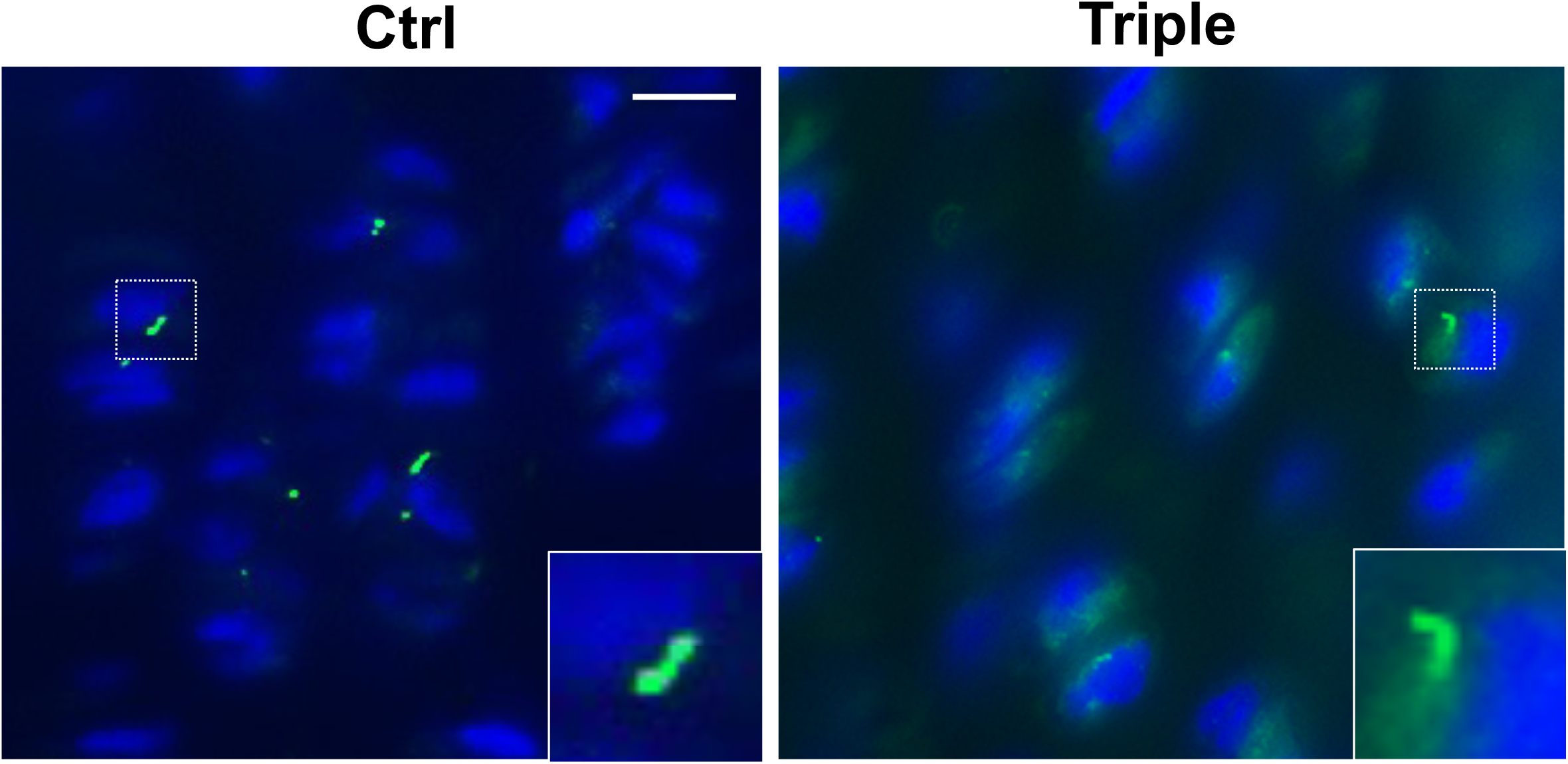
Cilia length is unaltered in *Thm2^-/-^; Thm1^aln/+^* cells. Immunostaining of P14 tibial growth plates for acetylated *α*-tubulin (green). Scale bar – 10 µm.

### Reducing Gli2 dosage in Thm2-null mice results in small phenotype

To determine whether Hh signaling contributes to the skeletal phenotype, we introduced a null allele of *Gli2* into the *Thm2; Thm1* colony. *Gli2-/-* mice show skeletal defects and perinatal lethality, but the *Gli2+/-* genotype does not result in a phenotype^45^. Yet the loss of one allele of *Gli2* in *triple; Gli2+/-* mice further decreased body weight and crown-to-rump length relative to triple allele mutant mice (Figure 9). Additionally, the loss of one allele of *Gli2* in *Thm2-/-; Gli2+/-* mice resulted in a small phenotype, with reduced body weight and crown-to-rump length relative to control mice. Thus, reducing Hh signaling uncovers a role for *Thm2* in postnatal skeletal growth. Further, *Gli2* deficiency demonstrates that *Thm2,* together with *Thm1,* positively regulates the Hh pathway in postnatal skeletal growth.

**Figure 9.**
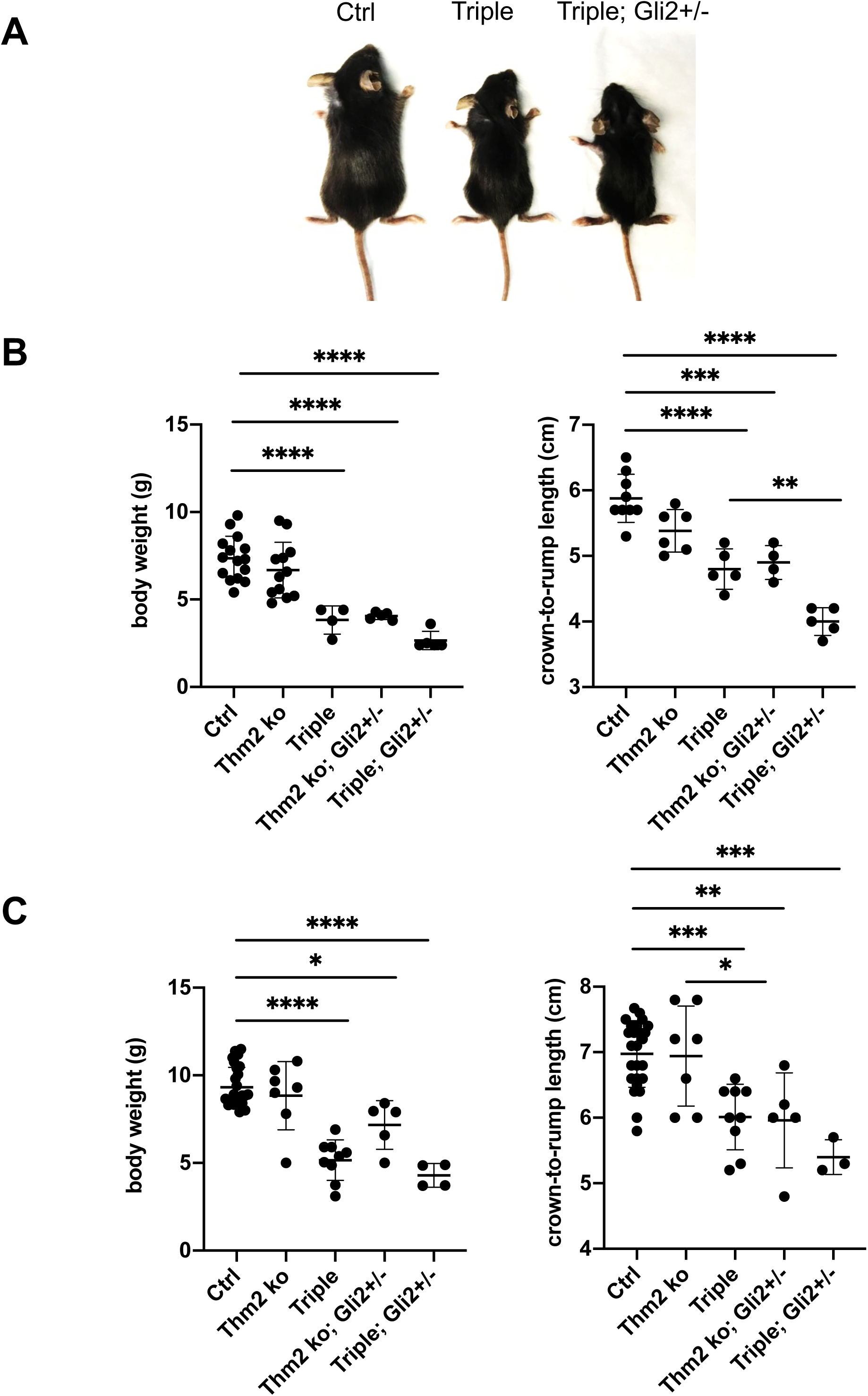
*Gli2* deficiency causes small phenotype in *Thm2* ko mice and exacerbates small phenotype in *Thm2^-/-^; Thm1^aln/+^* mice. A) Images of P14 littermates. B) Body weight and crown-to-rump lengths at P14 and C) at P21. Graphs represent mean ± standard deviation. Statistical significance was determined using one-way ANOVA and Tukey’s test. ****p≤0.0001

## Discussion

These studies establish *Thm2* as a ciliary protein. While depletion of *Thm2* results in normal cilia length, the loss of *Thm2,* together with either loss of one allele of *Thm1* or of *Gli2* causes abnormal postnatal skeletal development. These data increase our knowledge of the ciliary proteins and genetic interactions that regulate skeletal development. Additionally, the *Thm2^-/-^; Thm1^aln/+^* mouse is the first global *Ift*-null mouse surviving embryogenesis and exhibiting postnatal skeletal abnormalities.

Mutations in *THM1* have been identified in patients with Jeune Syndrome (JATD) ^27^. Our data show that *Thm2^-/-^; Thm1^aln/+^* mice model multiple aspects of Jeune Syndrome, including shortened long bones, micrognathia, and narrow ribcage. The most consequential abnormality of JATD is narrowing of the thoracic cavity, often resulting in respiratory insufficiency^38^. Interestingly, triple allele mutant mice show increased lung weight/body weight ratios, despite having reduced lung weight. This suggests that the overall body weight, and likely the skeleton, is smaller than normal to accommodate lung growth. Jeune Syndrome patients also exhibit varying degrees of severity^46^. Approximately 60-80% of patients die during the neonatal-infantile period of life^38^, while some patients are able to survive to adulthood^46^. Triple allele mutant mice also show varying degrees of severity, with a proportion of mutants not surviving to weaning. Our study also shows that by reducing the level of *Gli2*, triple allele mutants have an exacerbated phenotype and *Thm2* ko mice manifest a small phenotype. Moreover, genetic background plays a role in the small phenotype, since the phenotype was apparent only once the colony was backcrossed from a mixed FVB/C57BL6/J strain background to C57BL6/J over 5 generations. Thus, these genetic crosses model the varying severity seen in the human disease.

In the growth plate, Parathyroid hormone-related protein (PTHrP) is expressed from the perichondral cells and chondrocytes where it is able to promote proliferation of chondrocytes within the growth plate^43^. Once the level of PTHrP is below a certain threshold, pre-hypertrophic and early hypertrophic cells produce *Ihh* which affects a variety of skeletal processes, including increased chondrocyte proliferation within the growth plate, stimulation of PTHrP production, and differentiation of perichondral cells to osteoblasts of the bone collar^43^. Endochondral growth plates of *Ihh-*null mice exhibit a decrease in Hh signaling and proliferation, as well as a decrease in proliferation zone length at E14.5^47^. In contrast, conditional deletion of Suppressor of Fused (Sufu), a negative regulator of Hh signaling^48^, using a *Col2a1*-Cre recombinase resulted in increased *Ptch1,* a target of Hh signaling, accompanied by an increase in the proliferative zone and a decrease in the hypertrophic zone in the growth plate of E16.5 tibia explants^48^. An additional study revealed loss of *Sufu* in mice resulted in hypoplasia of the scapula, radius, ulna, and humerus at E14.5^49^. These studies indicate that in the absence of Hh signaling, chondrocytes undergo premature hypertrophy^50^, while inappropriate activation of Hh signaling causes chondrocytes to stay in the proliferation zone. Our analysis of P14 tibia growth plates in *Thm2^-/-^; Thm1^aln/+^* mice revealed an increased proliferation zone and a decreased hypertrophic zone, characteristic of increased Hh signaling. Yet deletion of one allele of *Gli2* exacerbated rather than rescued the triple allele mutant small phenotype. This suggests that there are likely other cells, such as osteocytes or osteoblasts, that play a more predominant role in the small phenotype. These data also suggest that in the causative cell type, the loss of *Thm2* together with one allele of *Thm1* reduces the ability to respond to the Hh signal. Similarly, we have found that triple allele mutant MEF have impaired ability to respond to the Hh agonist, SAG^44^.

*In vitro* knockdown of *Thm2* in 293T cells resulted in normal cilia lengths and the kidneys of P14 *Thm2^-/-^; Thm1^aln/+^* mice are normal. In contrast, loss of *Thm1* during late embryogenesis results in shortened primary cilia with a bulb-like tip and cystic kidneys at six weeks of age^51^. This suggests *Thm1* and *Thm2* share unique and redundant functions, and these likely vary by tissue type. Another example is polydactyly, which has not been observed in any of the *Thm2^-/-^; Thm1^aln/+^* mice, but manifests in several ciliopathies^38^, and in 100% of *Thm1^aln/aln^* mouse embryos^17, 32^. Polydactyly can be caused by disrupted GLI3A:GLI3R ratios along the anterior-posterior axis of the developing limb bud^52^. Since *Thm2* and *Thm1* show very similar expression patterns in E10.5 whole-mount mouse embryos^17^, the absence of polydactyly in *Thm2^-/-^; Thm1^aln/+^* mice may indicate divergent mechanisms of Hh regulation between *Thm2* and *Thm1*. *Thm1* loss disrupts GLI3 processing^17^, while *Thm2* likely does not.

Collectively, individuals with JATD show a wide spectrum of clinical abnormalities^46^. This may be attributed to genetic heterogeneity, since the mechanisms by which ciliary defects cause developmental defects can differ due to variations in function of individual ciliary proteins. For example, germline deletion of all reported *Ift* genes misregulates the Hh pathway, but not all affect ciliary structure^11, 53^. Moreover, whether a particular ciliary protein positively or negatively regulates the Hh pathway appears to be cell type- or context-dependent^19, 20, 54^.

Genetic analysis of individuals with JATD has revealed mutations in multiple cilia genes, including *THM1*^27^*, WDR19 (IFT144)*^25^*, DYNC2H1*^55, 56^*, IFT140*^26^, and *IFT80*^28, 57^. However in some patients with JATD, sequencing analysis of known genes did not uncover mutations, indicating additional loci contribute to the disease^38^. Our data establish that THM2 is a ciliary protein, and that *Thm2* interacts with *Thm1* or *Gli2* to positively regulate Hh signaling in skeletogenesis. These studies highlight important genic interactions during postnatal skeletal development and a novel locus to query in patients with JATD.

## Materials and Methods

### Generation of THM2 knockdown cell line

Lentiviral transfection was used to generate a 293T *Thm2* knockdown cell line. Briefly, viral particles were created by transfecting three plasmids, 4.2µg vector (*Thm2* shRNA sequence GACTTTGATTAATTACTAT), 7.4 µg delta 8.2 and 0.4 µg VSVG into 293T packaging cells using the Fugene transfection reagent (Promega, E2691) and incubated at room temperature for 20 minutes. The mixture was then incubated with 293T cells for 48 hours, after which supernatant was collected and filtered to obtain viral particles. Media was then placed onto 293T cells for 4 hours before media was changed. To obtain cells that had integrated the construct, cells were selected with 1µg/ml puromycin. Cells were seeded sparsely to allow growth and selection of cell clones using cloning disks immersed in Trypsin. Clones were then expanded.

### Western Blot

293T cells and mouse tissues were collected and protein was extracted using passive lysis buffer (Promega, E1941) containing protease inhibitors (Thermo Scientific, 88669). Briefly, samples were resuspended and mixed in lysis buffer for 15 minutes (293T cells) or resuspended in lysis buffer and homogenized (tissues). Cells and tissues were then pelleted and supernatants were collected. Sample lysates were prepared and loaded onto a 4-20% SDS-gel (BioRad, 456-8094). Gels were run for approximately 3.5 hours at 90V to allow for adequate separation between bands. Membranes were incubated with THM2 antibody using a Western blot procedure as described^17^. THM2 antibody was targeted to N’ Cys-RRQNYETAINLYHQVLEK, 963-980aa, in exon 22. (Proteintech S4132-2, 1:5,000).

### Immunofluorescence Staining

Retinal pigment epithelial (RPE) or human embryonic kidney (293T) cells were seeded on Poly-L-Lysine coated coverslips, grown, serum starved overnight, then fixed in 4% PFA in PBS containing 0.1% Triton x-100 for 10 minutes at room temperature. Fixed RPE or 293T cells were blocked in 2% Bovine Serum Albumin (Sigma, A9647) overnight at 4°C. Antibodies against acetylated α-tubulin (Sigma T6557, 1:4,000) and THM2 (Custom-made Antibody, Proteintech S4132-2, 1:50) were incubated with cells in 2% BSA overnight. Cells were incubated with secondary antibody (Life Technologies, A11005 and A11008, 1:500) for 1 hour at room temperature, inverted, and mounted on slides using fluoromount with DAPI (Electron Microscopy Services 17984-24). RPE cells were imaged on a Leica confocal microscope (Leica TCS SPE) while 293T cells were imaged on a light microscope (Nikon 80i) attached to a camera (Nikon DS-Fi1).

### Generation of THM2 knockout mouse

*Thm2* knockout-first C57BL/6J embryonic stem cells were obtained from the NIH Knockout Mouse Project (KOMP) Repository (www.komp.org). These embryonic stem cells were confirmed to contain the correct *Thm2* knockout construct exhibiting evidence of homologous recombination and correct karyotype, and subsequently, were injected into C57BL/6J-*Tyr^c-2J^* (albino) blastocysts and implanted into female mice by Dr. Melissa Larson at the University of Kansas Medical Center Transgenic and Gene Targeting Facility. Resulting male chimeras were mated to multiple females. Genomic DNA from tails of resulting pups with black coat color was genotyped by polymerase chain reaction (PCR) and PCR amplicons were sequenced to confirm presence of the construct in the DNA. The KOMP allele contains a LacZ neo construct flanked by flippase recombinase (FLP) recombinase target (FRT) sites. Mice containing the KOMP construct were mated to a mouse carrying a FLP recombinase (Jackson Laboratories, 009086) to excise the LacZ-Neo cassette. Resulting mice were then mated to mice expressing cytomegalovirus (CMV) Cre recombinase (Jackson Laboratories, 006064) to excise exon 6, which created a premature stop codon, and *Thm2* null allele.

### Genotyping of Thm2 knockout mouse

Genotyping primers were designed flanking the lox-P sites surrounding exon 6. Primer sequences used include: *Thm2-*KO*-*F-5’ CAG ATA TCT CCC CAC TTG TTA ACG 3’, and *Thm2-*KO*-*R-5’ GTG TCA GAT ACC CTG GAA CCA GAG 3’. With these primers, a WT band is amplified at a size of 1,086bp while a knockout band is amplified at a size of approximately 400bp. At times, the PCR reaction favors the amplification of the knockout band, making the WT band difficult to see. Therefore, to confirm the presence or absence of a WT allele, an additional PCR reaction is performed using primers within exon 6, *Thm2-*WT*-*F-5’AAC TTC CTG CCC GCT TTA GT 3’, and *Thm2-*WT*-*R-5’ GTG TCA GAT ACC CTG GAA CCA GAG 3’. This PCR reaction results in a WT amplicon of approximately 461bp, indicating the presence of a WT allele. PCR products are run on a 0.7% agarose gel for the detection of the recombined band and a 2% agarose gel for detection of the WT band.

### Generation of Thm2^-/-^;Thm1^aln/+^ mice

*Thm2^-/-^;Thm1^aln/+^* mice were generated by intercrossing *Thm2^-/+^* mice on a C57BL/6 background with *Thm1^aln/+^* mice on an FVB background. *Thm2^-/+^;Thm1^aln/+^* mice were backcrossed five generations onto a C57BL6/J background and then intercrossed to generate *Thm2^-/-^;Thm1^aln/+^* mice.

### Weight and Length Measurements

Total mouse body weight was measured in grams using a standard weighing scale. Crown to rump measurements in centimeters were taken manually using a standard ruler, measuring the length from the tip of the nose to the end of the rump. Total body weights and crown to rump measurements were recorded at P14 and P21.

### Skeletal Preparations

Alizarin red and alcian blue staining was performed using standard protocols (Kingsley Lab Protocol, Ryan Roundtree, Version 1.1, 3/31/2003). Briefly, P14 mice were eviscerated and fixed in 95% Ethanol for 1-2 days, and stained with alcian blue (Acros Organics 33864-99-2) for 14 days. After alcian blue staining, skeletons were fixed with 95% ethanol for an additional 1-2 days and cleared with potassium hydroxide for approximately 2-5 days, or until tissue was cleared. Skeletons were then stained with 1% alizarin red (Acros Organics 130-22-3) in potassium hydroxide for 1-2 days or until completely stained. Once staining was complete, skeletons were placed into glycerol for imaging and long-term storage. Skeletal preparations were then imaged (Leica M165C) and measured using Image J software.

### Microcomputed tomography and Faxitron

Whole mice and tibias were radiographed at 1X and 3X magnifications, respectfully, using a LX-60 DC12 Cabinet X-ray System (Faxitron) at 26kVP for 10 seconds. The images were analyzed in ImageJ to calculate the length of each mineralized caudal vertebrae.

Ethanol-fixed tibia from control and triple allele mutant mice were scanned in a Skyscan 1174 micro-computed tomography system (Bruker) at a resolution of 14 μ m3 with a voltage of 50kVp using a 0.5mm aluminum filter. Tibias were wrapped in gauze soaked with 70% ethanol and placed in a low density polyethylene tube for scanning. Camera pixel binning was not applied and the integration time was set to 3000 ms. The scan orbit was 180/360 degrees with a rotation step of 0.3 degrees. Reconstruction was carried out with a modified Feldkamp algorithm using the SkyScanTM NRecon software accelerated by GPU3. Gaussian smoothing, ring artifact reduction, and beam hardening correction were applied. Reconstructed images were analyzed by CTAn software. Irregular, anatomic regions of interest (ROIs) were selected for both trabecular and cortical bone analysis. For trabecular bone analysis, slices from proximal tibia starting at the distal end of the growth plate to the end of the trabeculae were analyzed for bone mineral density (BMD), bone volume fraction (BV/TV), bone surface per BV (BS/BV), trabecular thickness (Tb.Th), separation (Tb.Sp), and number (Tb.N). For cortical bone analysis, two sets of ROIs were analyzed for BMD, total cross-sectional area (Tt.Ar), cortical bone area (Ct.Ar), fraction (Ct.Ar/Tt.Ar), perimeter (Ct.Pm), and thickness (Ct.Th). The first was from mid-diaphysis and the second started at the end of the trabeculae at the proximal end of the tibia. To control for length differences, the same percentage of slices were analyzed in mutants as relative length for both trabecular and cortical analysis. The minimum and maximum thresholds were set to 45 and 95 for trabecular bone analysis and 55 and 255 for cortical bone analysis.

### Dissection, Embedding and Sectioning of Tibias and Soft Organs

Tibias of mice were dissected and placed in Cal-Ex, a decalcifying and fixing solution (Fisher C5511-1D) overnight. Kidney, heart and lung were dissected and weighed. Kidneys were fixed in Bouin’s fixative (Polysciences, Inc. 16045) overnight. Tibias and kidneys were placed into 70% ethanol. Tissues were processed and dehydrated through a series of ethanol washes and embedded in paraffin wax. Paraffin blocks were sectioned at 10μm and 7μm for tibias and soft organs, respectively.

### Hematoxylin and Eosin Staining

Tibia and kidney sections were stained using standard hematoxylin and eosin staining methods. Briefly, sections were de-paraffinized and rehydrated through a series of ethanol rinses. Sections were then stained with Hematoxylin (Sigma HHS32) and Eosin (Sigma HT110216) before being dehydrated and mounted with Permanent Mounting Media (Fisher, SP15-500). Sections were imaged using a light microscope (Nikon 80i) attached to a Nikon DS-Fi1 camera.

### Safranin O and Fast Green Staining

P14 tibia sections were stained using Safranin O and Fast Green Staining (American MasterTech, KTSFO). Briefly, sections were de-paraffinized and rinsed in 100% ethanol. Slides were stained with Weigert’s Hematoxylin, 0.2% Fast Green, and Safranin O in separate steps. After staining, sections were dehydrated in 100% Ethanol, cleared in Xylene and mounted using Permanent Mounting Media (Fisher SP15-500).

### Statistics

Unpaired, two-tailed t-tests for comparison of two groups and one-way ANOVA followed by Tukey’s test for comparison of multiple groups were used to evaluate statistical significance (p<0.05).

## Acknowledgements

We thank members of the KUMC Dept. of Anatomy and Cell Biology, the Jared Grantham Kidney Institute, and the UMKC Dept. of Oral and Craniofacial Sciences for helpful discussions. We thank Jing Huang of the KUMC Histology Core and Drs. Vivian and Larson of the KUMC Gene Targeting Institution Facility and acknowledge support of these cores (Intellectual and Developmental Disabilities Research Center NIH U54 HD090216; KU Cancer Center NIH P30 CA168524; COBRE NIH P30 GM122731). This work was also supported by a KUMC Biomedical Research Training Fellowship, a pilot grant from the Kansas City Consortium on Musculoskeletal Diseases, and the National Institutes of Health (P20 GM14936; R01DK103033).

## Contributions

BAA, WW, TSP, HM, LMS, DTJ, JW, EEB, and PVT performed experiments and analyzed data. BAA, JW, EEB and PVT designed research. BAA, HM, EEB and PVT wrote the paper.

---

**Figure S1. Cortical and trabecular properties of *Thm2^-/-^; Thm1^aln/+^* tibia.** A) Cortical bone area, total area, cortical area fraction (cortical bone area/total area), and cortical bone perimeter at the proximal end of the tibia B) and at mid-diaphysis. C) Trabecular number, trabecular separation, trabecular bone surface/bone volume, and trabecular volume/total bone volume. Bars represent mean ± standard deviation of n=2 ctrl and n=2 triple allele mutant mice.

**Figure S2. *Thm2^-/-^; Thm1^aln/+^* cilia length is not altered.** A) Immunostaining of MEF for ARL13B (green) and *γ*-tubulin (red). Scale bar - 5μm. B) Quantification of cilia length. Data points represent individual cilia. Graphs represent mean ± standard deviation. Statistical significance was determined using one-way ANOVA and Tukey’s test. *p*≤*0.05

**Movie S1. Microcomputed tomography video of P14 control tibia.**

**Movie S2. Microcomputed tomography video of P14 triple allele mutant tibia.**

